# Proteasome maturation factor UMP1 confers broad-spectrum disease resistance by modulating H_2_O_2_ accumulation in rice

**DOI:** 10.1101/2021.03.03.433750

**Authors:** Xiao-Hong Hu, Jing Fan, Jin-Long Wu, Shuai Shen, Jia-Xue He, Jie Liu, He Wang, Yong Zhu, Guo-Bang Li, Jing-Hao Zhao, Jie Xu, De-Qiang Li, Mei Pu, Zhi-Xue Zhao, Shi-Xin Zhou, Ji-Wei Zhang, Yan-Yan Huang, Yan Li, Yan-Li Lu, Fu Huang, Wen-Ming Wang

**Affiliations:** State Key Laboratory of Crop Gene Exploration and Utilization in Southwest China, Sichuan Agricultural University, Chengdu, 611130, China; Maize Research Institute, Sichuan Agricultural University, Chengdu, 611130, China; College of Agronomy, Sichuan Agricultural University, Chengdu, 611130, China

**Author notes:** These authors contributed equally to this work (Xiao-Hong Hu, Jing Fan).

## Abstract

Crops with broad-spectrum resistance (BSR) to diseases are highly desirable in agricultural production. Identification of BSR loci and dissection of the underlying mechanisms are fundamental for crop resistance breeding. Here, we describe the identification and characterization of a rice *UMP1* allele, which confers race-nonspecific BSR against blast pathogen *Magnaporthe oryzae*. *OsUMP1* encodes a proteasome maturation factor that contributes to 26S proteasome abundance and activity in rice. Modulation of *OsUMP1* expression leads to proteome changes, particularly affects the amounts and activities of H_2_O_2_-degrading enzymes. Consequently, H_2_O_2_ accumulation and disease resistance are enhanced in *OsUMP1-*overexpressing rice but reduced in loss-of-function mutants. Elevation of *OsUMP1* expression also promotes rice resistance to foliar pathogens *Rhizoctonia solani* and *Xanthomonas oryzae* pv. *oryzae* and a floral pathogen *Ustilaginoidea virens* without observable yield penalty. These results indicate a BSR pathway linking the proteasome machinery and H_2_O_2_ homeostasis, and provide a candidate gene for balancing BSR and yield traits in rice breeding.

**One Sentence Summary:** A natural allele of rice UMP1 promotes resistance to multiple pathogens by boosting H_2_O_2_ accumulation.

## INTRODUCTION

Crops are suffered from many plant diseases caused by fungi, bacteria, oomycetes, nematodes, and viruses, resulting in great reduction of yield and quality. Application of pesticides for controlling diseases can impose environmental and health threats. Development of resistant crop cultivars is considered as the most economical and eco-friendly way to manage diseases. Crops with broad-spectrum resistance (BSR) against most strains/races of a pathogen or more than one pathogen species (Li et al., 2020) are highly desirable by plant breeders and producers. Breeding for elite cultivars with BSR would benefit greatly from the identification and characterization of superior BSR loci.

A list of BSR genes have been isolated in plants and can be classified into seven subtypes, including membrane-associated pattern recognition receptor genes, nucleotide-binding and leucine-rich repeat receptor genes, defense-signaling genes, pathogenesis-related genes, mutated susceptibility genes, quantitative trait loci, and nonhost-resistance genes (Li et al., 2020). Some of them have been mechanistically characterized. For instance, *Lr67/Yr46*, encoding a hexose transporter, can modulate hexose transport to promote resistance against powdery mildew, leaf rust, stem rust, and stripe rust pathogens in wheat (Moore et al., 2015). *Fusarium head blight 7* (*Fhb7*) confers resistance to FHB and crown rot in different wheat backgrounds through a mechanism of detoxifying trichothecenes by de-epoxidation (Wang et al., 2020). The Arabidopsis *RPW8* loci confers resistance to both *Erysiphe cichoracearum* and *Hyaloperonospora parasitica* by inducing salicylic acid-dependent basal defenses (Xiao et al., 2001; Wang et al., 2007). Interfamily transfer of *RPW8.1* to rice can boost plant pattern-triggered immunity (PTI) and resistance to rice blast and bacterial blight (Li et al., 2018).

As a staple crop, rice has been threatened by multiple diseases, mainly including rice blast (caused by *Magnaporthe oryzae*), sheath blight (caused by *Rhizoctonia solani*), and bacterial blight (caused by *Xanthomonas oryzae* pv. *oryzae*, *Xoo*) (Liu et al., 2014; Yin et al., 2018). Rice false smut, caused by the flower-infecting pathogen *Ustilaginoidea virens*, has recently emerged as one of the most notorious diseases in rice (Sun et al., 2020). Advances have been made to identify rice BSR genes (Li et al., 2020), such as *pi21*, *PigmR*, *bsr-d1*, *bsr-k1*, *xa13,* and *Xa21* (Song et al., 1995; Chu et al., 2006; Fukuoka et al., 2009; Deng et al., 2017; Li et al., 2017; Gao et al., 2018; Zhou et al., 2018). *PigmR* isolated from a rice cultivar Gumei4 encodes a nucleotide-binding leucine-rich repeat receptor and provides durable and BSR to diverse isolates of *M. oryzae* (Deng et al., 2017). Digu-originated *bsr-d1* is a natural allele of a C_2_H_2_-family transcription factor, and contributes to inhibition of H_2_O_2_ degradation leading to enhanced resistance to a diverse panel of *M. oryzae* isolates (Li et al., 2017). *Xa13* encodes a sugar transporter (OsSWEET11) acting as a susceptibility factor to support pathogen growth. Loss-of-function of *Xa13* enhances rice resistance to both *Xoo* and *R. solani* (Chu et al., 2006; Gao et al., 2018). So far, however, no resistance genes have been identified to function against the floral pathogen *U. virens* (Sun et al., 2020).

We previously developed an elite hybrid rice restorer line Yahui2115 (R2115), which shows BSR to rice blast (Shi et al., 2015) and effective field resistance to rice false smut (the highest disease index = 1.7% from more than 10 years of field evaluations). Thus, R2115 can be served as an excellent resource for exploring novel genes conferring BSR. In this study, we identified a natural allele of a proteasome maturation factor *OsUMP1* that promoted resistance to both rice blast and rice false smut, as well as bacterial blight and sheath blight. *OsUMP1* may contribute to BSR through regulating proteasome activity and inhibiting H_2_O_2_ degradation, representing a novel mechanism of rice disease resistance.

## RESULTS

### Identification of R2115-specific genes responsive to *Magnaporthe oryzae* infection

To explore BSR genes in R2115, we performed comparative transcriptome analysis on *M. oryzae*-challenged R2115, susceptible rice Lijiangxin Tuan Heigu (LTH) and three of its derived monogenic resistant lines, namely IRBLZ5-CA, IRBLZ9-W, and IRBLKM-TS harboring race-specific blast resistance gene *Pi2*, *Pi9* and *Pikm*, respectively. We reasoned that genes specifically induced in R2115 but not in IRBLZ5-CA, IRBLZ9-W, and IRBLKM-TS, may contribute to BSR. At 6 days post inoculation (dpi) of *M. oryzae*, seedling leaves of LTH displayed susceptible lesions, whereas leaves of R2115, IRBLZ5-CA, IRBLZ9-W and IRBLKM-TS showed no lesions at all, confirming their resistance to examined *M. oryzae* isolates (Fig. 1A). Accordingly, the expressions of defense-related genes were induced to higher levels in the resistant lines, compared to the susceptible LTH (Fig. 1B).

**Figure 1.**
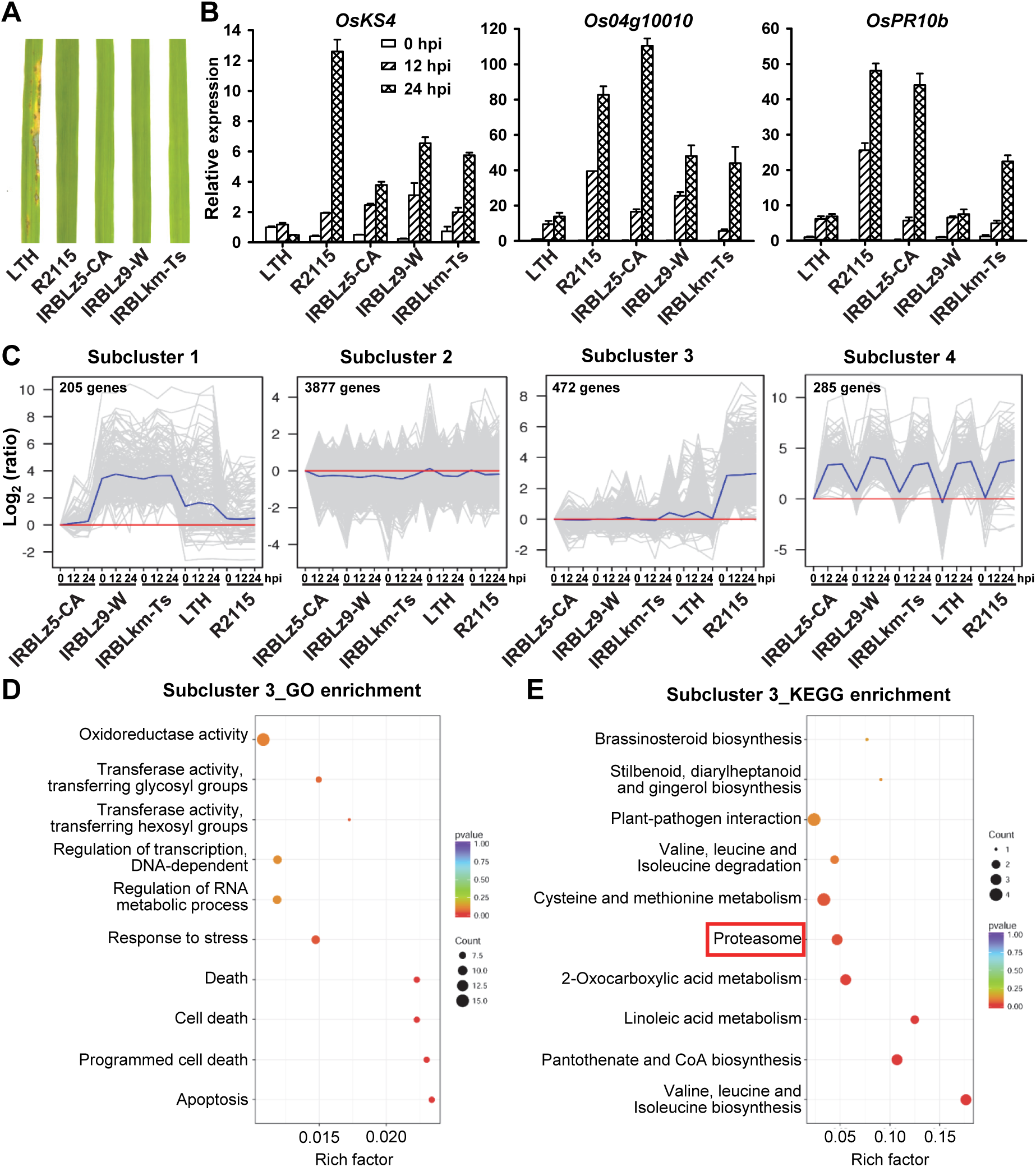
Identification of R2115-specific genes responsive to *Magnaporthe oryzae* infection. **A,** Disease phenotype of leaves from indicated lines at seven days post inoculation of *M. oryzae*. **B,** Expression analysis of indicated defense-related genes at indicated time points of *M. oryzae* infection. *OsUbi* was used as the reference gene. The expression of indicated genes in LTH-0 hpi was set as the control. Data are means ± SD of three biological replicates. **C,** Expression patterns of Subcluster 1-4 differentially expressed genes (DEGs). Comparative transcriptome analysis was performed on indicated rice lines at multiple time points of *M. oryzae* infection. All DEGs were subjected to Hierarchical-clustering analysis, which generated 10 subclusters. Subclusters 5-10 were presented in Supplemental Figure S1. **D** and **E,** GO and KEGG enrichment analysis of DEGs in Subcluster 3. Top 10 terms were presented. The term of interest was marked by a red rectangle. Rich factor refers to the ratio of DEG number against annotated gene number in background genome for a certain term or pathway.

Subsequently, leaf samples collected at 0, 12, 24 h post inoculation (hpi) were used for RNA-Seq analysis. Totally, 30 libraries were sequenced and each generated 50 million reads in average (Supplemental Table S1). Differentially expressed gene (DEG) analysis revealed 16924 genes showing significantly different expression levels between any two samples (≥ 2-fold change and false discovery rate < 0.05). These DEGs were grouped into 10 subclusters according to their expression profiles (Fig. 1C; Supplemental Fig. S1; Supplemental Table S2). Interestingly, 472 DEGs of Subcluster 3 displayed *M. oryzae*-inducible and higher expression in R2115 than in LTH, IRBLZ5-CA, IRBLZ9-W, and IRBLKM-TS (Fig. 1C). This gene set was selected as BSR candidate for further GO and KEGG enrichment analyses. Subcluster 3 DEGs were enriched in GO/KEGG terms such as programmed cell death and proteasome (Fig. 1D, E; Supplemental Table S3, S4). Among DEGs involved in these pathways, a gene (Os03g0583700) encoding a putative proteasome maturation factor UMP1 was markedly expressed with a pathogen-inducible pattern in R2115 (Fig. 2A; Supplemental Table S5). Three *UMP1* homologs harbored in rice genome include Os03g0583700, Os09g0314900, and Os02g0800100, which displayed diverse expression profiles in our transcriptome data (Supplemental Fig. S2). Os09g0314900 was expressed at low or undetectable levels, while Os02g0800100 was constitutively expressed in all examined samples (Supplemental Fig. S2). Therefore, Os03g0583700 (hereafter *OsUMP1*) was selected for further functional identification.

**Figure 2.**
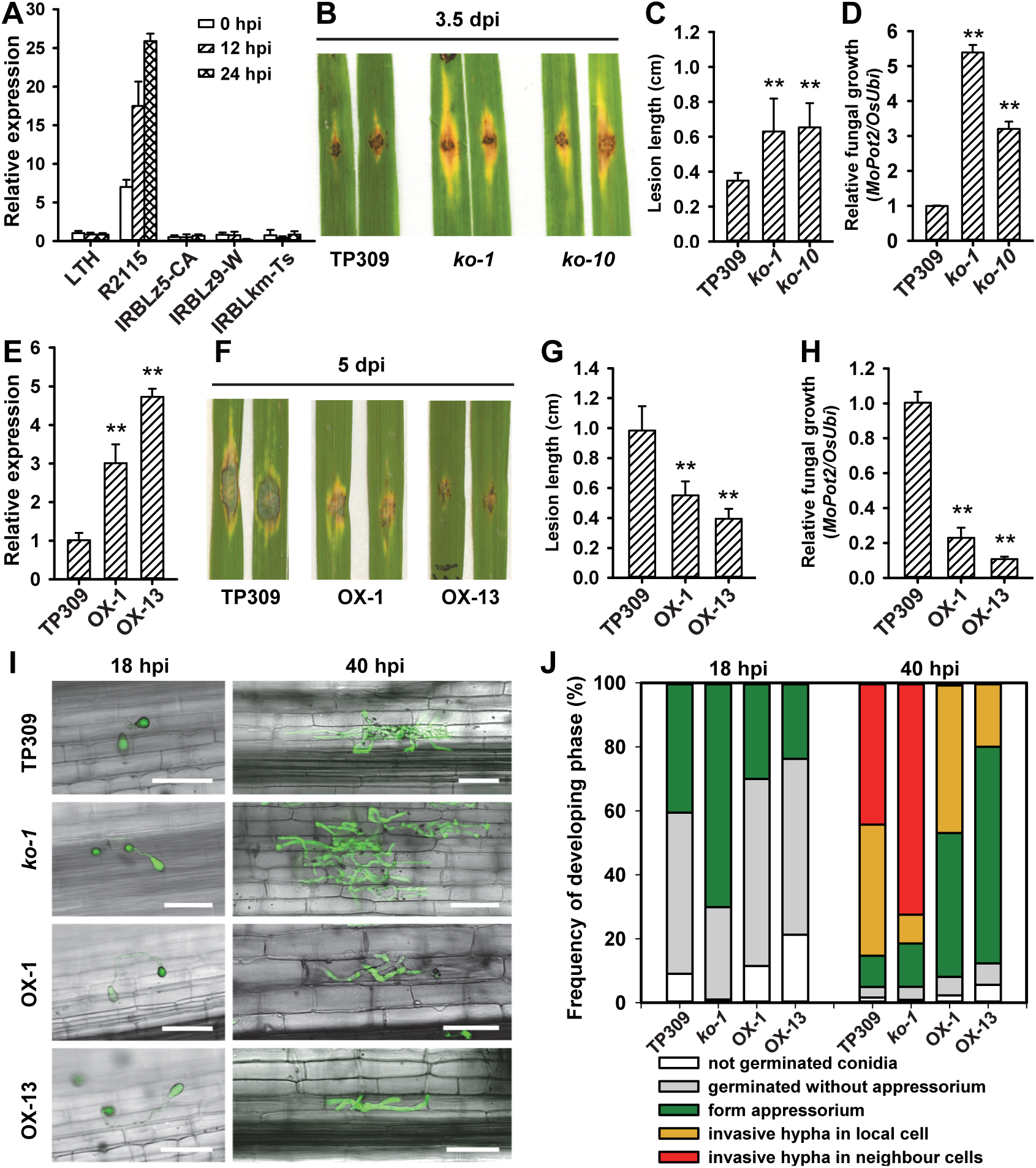
*OsUMP1* positively regulates rice resistance to *Magnaporthe oryzae*. **A,** RT-qPCR analysis of *OsUMP1* in indicated rice lines at indicated time points of *M. oryzae* infection. Os*Ubi* was used as the reference gene. The expression of *OsUMP1* in LTH-0 hpi was set as the control. Values are means ± SD of three biological replicates. **B,** Disease phenotype of *OsUMP1* knockout lines *ko-1* and *ko-10* at 3.5 days post inoculation (dpi) of *M. oryzae* GZ8. **C** and **D,** Quantification of lesion length (C) and relative fungal growth (D). Values are means ± SD of six replicates. Asterisks indicate significant differences between TP309 and knockout lines as detected by Student’s *t*-test (** *P* < 0.01). **E,** Expression level of *OsUMP1* in *OsUMP1^R2115^*-overexpressing transgenic plants OX-1 and OX-13. The expression level of *OsUMP1* was determined by RT-qPCR using *OsUbi* as the reference gene and TP309 as the control. Values are means ± SD of three replicates. Asterisks indicate significant differences between TP309 and OX lines as determined by Student’s *t*-test (** *P* < 0.01). **F,** Disease phenotype of *OsUMP1*-overexpressing lines at 5 dpi of GZ8. **G** and **H,** Quantification of lesion length (G) and relative fungal growth (H). Values are means ± SD of six replicates. Significant difference between TP309 and OX lines was determined by Student’s *t*-test (** *P* < 0.01). **I,** Representative confocal images of rice sheath cells infected by GZ8 at indicated time points. Scale bars, 50 μm. **J,** Frequency of developing phase of GZ8 in the sheath cells of indicated rice lines at 18 and 40 hpi. More than 150 spores at each time point in each line were analyzed.

### *OsUMP1* contributes to rice resistance to *M*. *oryzae*

To determine the functions of *OsUMP1*, we first intended to knockout it in a *M. oryzae*-compatible rice accession Taipei 309 (TP309) rather than in R2115, as virulent *M. oryzae* isolates to R2115 have not been identified. Based on CRISPR-Cas9 technology, we designed a sgRNA to target the first exon of *OsUMP1* and obtained mutants with either 1-bp insertion or 2-bp deletion at the target site (Supplemental Fig. S3A), introducing early stop codons and resulting in truncated proteins (Supplemental Fig. S3B). *osump1* mutant lines were then inoculated with *M. oryzae* strain GZ8. Compared to the wild-type TP309 and/or heterozygous mutant plants, *osump1* mutants consistently showed larger blast lesions and higher amounts of *M. oryzae* biomass in both T0 (Supplemental Fig. S3C-E) and T2 (Fig. 2B-D) generations. Next, we introduced the *OsUMP1* allele (native promoter plus gene region) from R2115 into TP309 and generated multiple transgenic lines with elevated expression levels of *OsUMP1*, which were associated with their resistance phenotype to *M. oryzae* (Supplemental Fig. S3F, G). Two representative transgenic lines OX-1 and OX-13 showing high resistance level were used for subsequent experiments (Fig. 2E). Upon infection with *M. oryzae* GZ8, OX-1 and OX-13 displayed much smaller lesions and supported less *M. oryzae* fungal biomass than TP309, indicating enhanced resistance to *M. oryzae* (Fig. 2G-H). Assessment of segregating progenies confirmed that this resistance phenotype co-segregated with the transgene and *OsUMP1* expression level (Supplemental Fig. 3H-K). Moreover, infection progression analysis showed that *osump1* mutant line *ko-1* displayed elevated rate of appressorium infection at 18 hpi and increased invasive hyphae progression at 40 hpi with GZ8 in sheath cells; by contrast, OX-1 and OX-13 showed delayed infection progress of GZ8, compared to the TP309 control (Fig. 2I-J). Taken together, these data indicate that *OsUMP1* contributes to rice resistance against *M. oryzae*.

**Figure 3.**
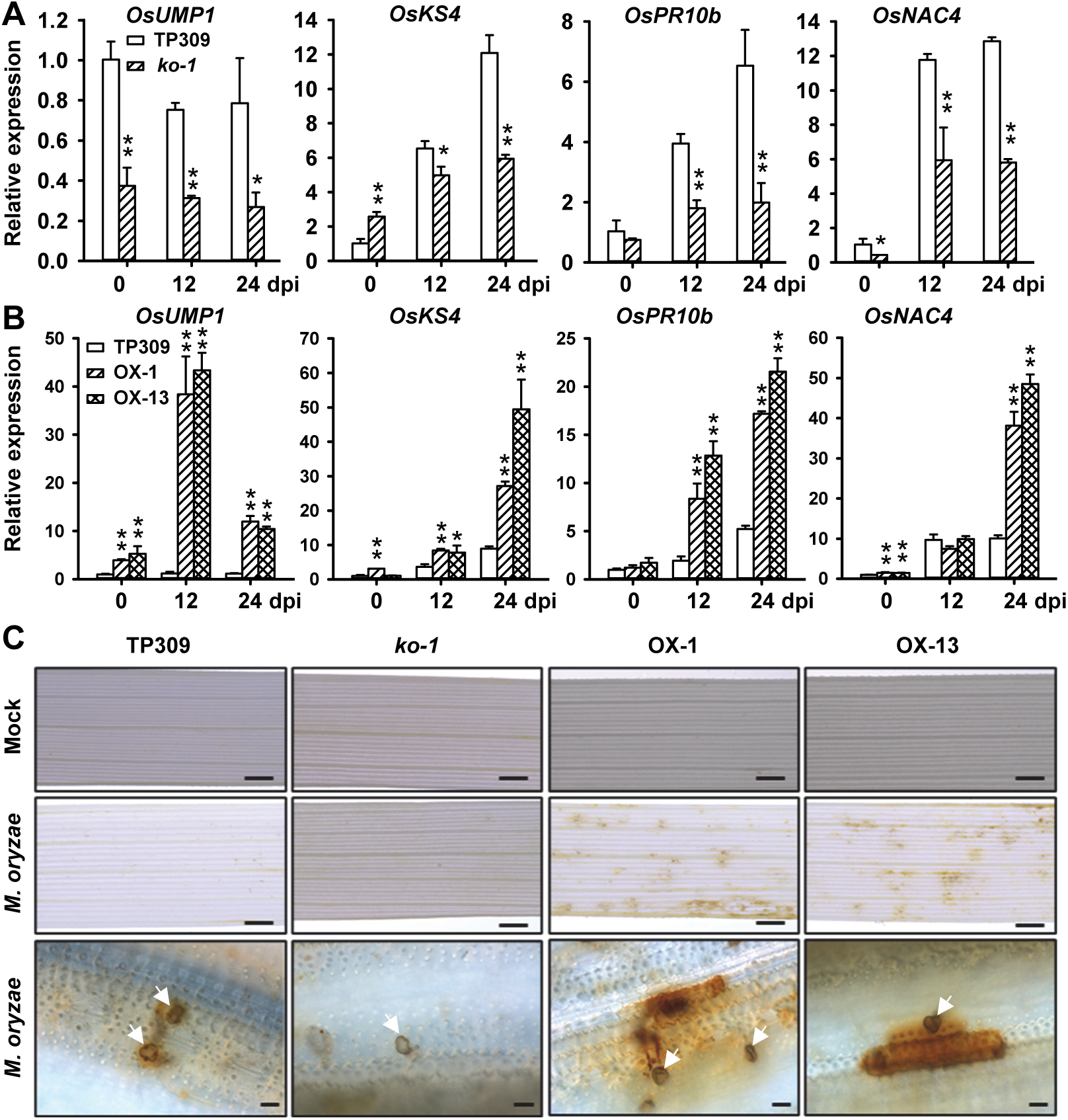
*OsUMP1* regulates rice immune responses. **A** and **B,** Expression analysis of *OsUMP1* and defense-related genes in leaves of indicated rice lines after spray-inoculation of GZ8. *OsUbi* was used as the reference gene. The expression of indicated genes in TP309-0 hpi was set as the control. Data are means ± SD of three biological replicates. Significant difference between TP309 and transgenic lines was determined by Student’s *t*-test (* *P* < 0.05, ** *P* < 0.01). **C,** GZ8-challenged leaves of indicated rice lines were stained with DAB at 48 hpi. Mock-inoculated leaves were served as the control. H_2_O_2_ accumulation was indicated by dark brown color. White arrows indicate appressoria of *M. oryzae*. Size bars =1 mm (top and middle panels), 10 μm (bottom panel).

### *OsUMP1* regulates rice immune responses

To understand how *OsUMP1* might confer resistance to *M. oryzae*, we examined the defense responses of *OsUMP1* transgenic plants inoculated with *M. oryzae*. Following pathogen infection, the expression of *OsUMP1* was induced to a high level in OX-1 and OX-13, but remained non-induced at a low level in TP309 and much lower in knockout mutant *ko-1* (Fig. 3A-B). Consistently, the expressions of defense-related genes including *OsKS4*, *OsPR10b*, and *OsNAC4* were less induced in *ko-1*, while induced to much higher levels in OX-1 and OX-13, compared to TP309 (Fig. 3A-B). At 48 hpi, inoculated rice leaves were stained with DAB for monitoring H_2_O_2_ accumulation. TP309 and mutant *ko-1* leaves accumulated low or no detectable levels of H_2_O_2_ around infection sites of *M. oryzae*, whilst OX-1 and OX-13 leaves were stained to dark brown at infection sites, indicating high abundance of H_2_O_2_ accumulation (Fig. 3C). These findings indicate that overexpression of *OsUMP1* boosts rice immune responses against *M. oryzae*, and loss-of-function of *OsUMP1* attenuated rice immunity.

### *OsUMP1* regulates proteasome activity and H_2_O_2_ accumulation in rice

The *Saccharomyces cerevisiae* homolog *Ump1p* is involved in regulation of proteasome maturation and assembly, leading to formation of active proteasome (Ramos et al., 1998). Thus, we tested whether *OsUMP1* genetically contributes to the accumulation and activity of 26S proteasome in rice. Using an ELISA approach, we first measured the amounts of 26S proteasome in *OsUMP1* overexpression and knockout plants. The abundance of 26S proteasome accumulated significantly higher in OX-1 and OX-13 than in TP309 before/after *M. oryzae* infection (Fig. 4A); while the 26S proteasome amounts were decreased in *ko-1* only upon *M. oryzae* infection, compared to the TP309 control (Fig. 4B). Using a fluorogenic peptide Suc-LLVY-AMC as the substrate (Üstün and Börnke, 2017), we then determined the 26S proteasome activity in above rice plants infected with *M. oryzae*. The 26S proteasome activity was significantly higher in OX-1 and OX-13 but lower in *ko-1* than in TP309 before/after *M. oryzae* infection (Fig. 4A-B). We also observed that the differences of proteasome activity among tested rice lines tended to be larger than those of proteasome abundance (Fig. 4A-B), indicating that *OsUMP1* contributes greatly to proteasome activity.

**Figure 4.**
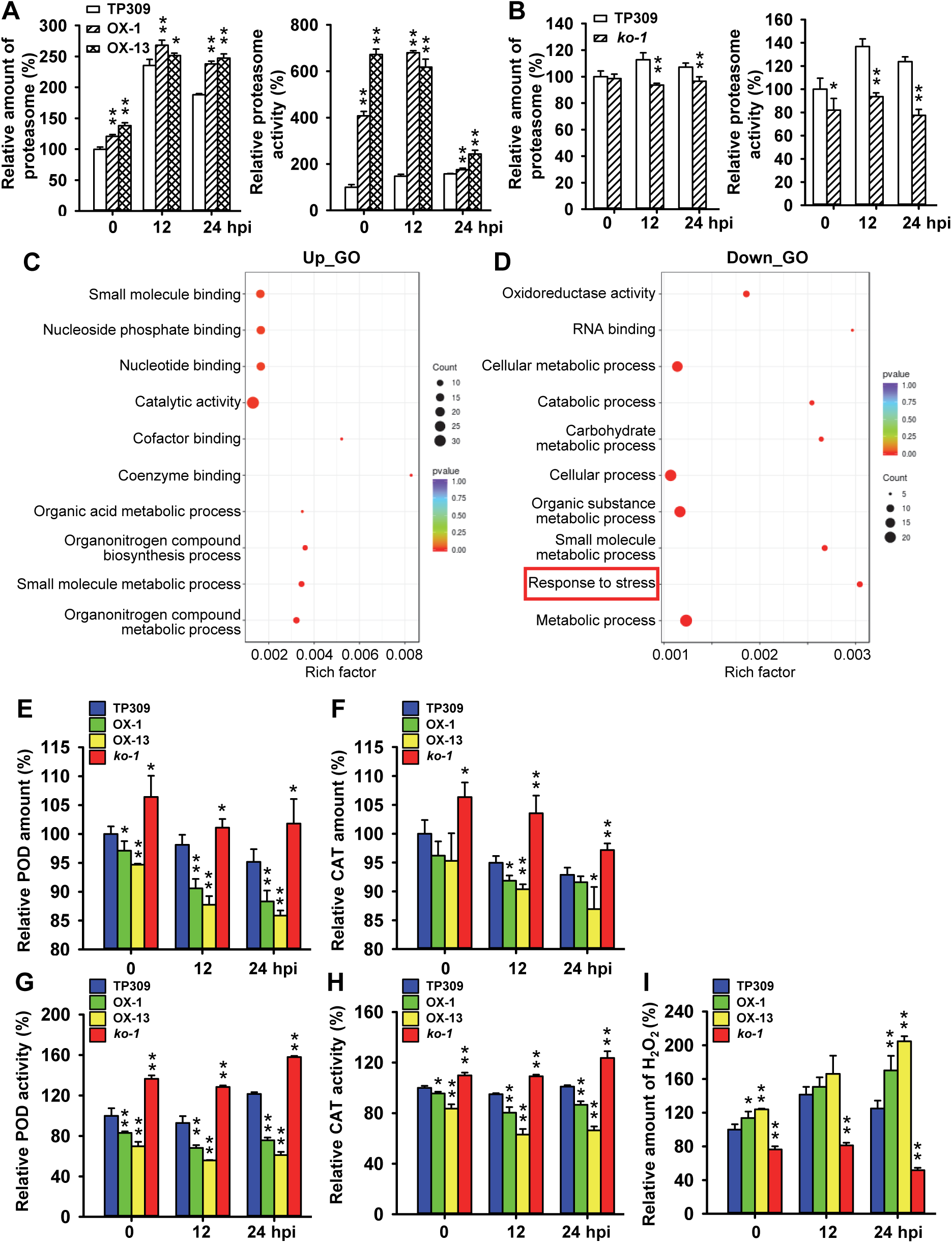
*OsUMP1* regulates abundance and activity of 26S proteasome and H_2_O_2_ accumulation in rice. **A** and **B,** The 26S proteasome amount (A) and activity (B) in leaves of indicated rice lines after spray-inoculation of GZ8. **C** and **D,** GO enrichment analysis of differentially accumulated proteins (DAPs) between TP309 and *OsUMP1*-overexpressing line OX-1. Leaves from four-leaf-stage seedlings were used for iTRAQ proteomic analysis. Top 10 enriched GO terms were presented. The term of interest was marked by a red rectangle. Rich factor refers to the ratio of DAP number against annotated protein number in background genome for a certain term. **E** and **F,** ELISA assay determining the abundance of POD (E) and CAT (F) in leaves of indicated rice lines after spray-inoculation of GZ8. **G** and **H,** Enzyme activities of POD (G) and CAT (H) in leaves of indicated rice lines after spray-inoculation of GZ8. I, Quantification of H_2_O_2_ in leaves of indicated rice lines after spray-inoculation of GZ8. Data are means ± SD of three biological replicates. Significant difference between TP309 and transgenic lines was determined by Student’s *t*-test (* *P* < 0.05, ** *P* < 0.01).

Since proteasome is a vital machinery for global protein degradation in plant cells, we monitored the proteome changes caused by alteration of *OsUMP1* expression with an iTRAQ approach. Comparative proteomic analysis revealed that, out of 2767 identified proteins, 149 proteins accumulated at higher levels and 103 proteins accumulated at lower levels in OX-1 than in TP309 (fold change ≥1.2 and *P* value <0.05; Supplemental Table S6). Gene ontology enrichment analysis demonstrated that up-regulated proteins were enriched in 26 biological processes/functional activity/cellular component, such as “organonitrogen compound metabolic process” and “small molecule binding” (Fig. 4C; Supplemental Table S7). By contrast, the most enriched terms for down-regulated proteins were “metabolic process” and “response to stress” (Fig. 4D; Supplemental Table S7). We further looked into the category of “response to stress”, and found that the protein levels of three ascorbate peroxidases (B9FV80, A3A7Y3, and B8AU10) and a catalase (A0A0R7FMX6) decreased to 69.7%-81.6% in OX-1 compared to those in TP309 (Supplemental Table S6).

We intended to validate the proteome data by raising antibodies for several differentially accumulated proteins, such as B9FV80, A3A7Y3, A0A0R7FMX6, and Q9XHY3. However, we failed to obtain antibodies against those candidates. We next sought out to examine POD and CAT abundance in *OsUMP1* transgenic lines and TP309 by an ELISA approach, and observed that both POD and CAT proteins accumulated less in OX-1 and OX-13 but more in *ko-1*, compared to the TP309 control before/after *M. oryzae* infection (Fig. 4E, F). We also determined the enzymatic activities of POD and CAT in above plants, and found that POD and CAT activities showed similar trends to their abundance (Fig. 4G, H). These results demonstrate disturbation of H_2_O_2_-degrading enzymes resulted from changes of *OsUMP1* expression and proteasome activity.

We next quantified H_2_O_2_ accumulation in *OsUMP1* transgenic lines and TP309 before/after *M. oryzae* infection. The basal abundance of H_2_O_2_ displayed approximately 20% difference between TP309 and transgenic lines (Fig. 4I). At 24 hpi with *M. oryzae*, the differences of H_2_O_2_ accumulation became larger among TP309 and transgenic lines, i.e., 36%-64% increases in *OsUMP1-*overexpressing lines and approximately 59% decreases in the knockout line, compared to the TP309 control (Fig. 4I). Taken together, our data suggest that overexpression of *OsUMP1* leads to reduction of H_2_O_2_-degrading enzymes and in turn enhances H_2_O_2_ accumulation to promote disease resistance, and vice versa.

### *OsUMP1^R2115^* is a rare allele in rice populations and has potential value in application

To investigate the natural variations in *OsUMP1*, we analyzed its DNA polymorphism among over 5000 rice accessions with MBKbase (http://mbkbase.org/rice) (Peng et al., 2020). Using default parameters, we identified 157 haplotypes for 2.5-kb promoter plus gene region of *OsUMP1* (Supplemental Table S8). There were 42 SNPs or InDels in the promoter region but only four SNPs or InDels in the coding region of *OsUMP1* (Fig. 5A; Supplemental Table S8). Little variations in the coding region of *OsUMP1* suggests functional conservation of OsUMP1 protein in rice. This notion was supported by that OsUMP1 protein sequence (particularly the C-terminus) was highly conserved among different plant species, such as wheat, maize, sorghum, and cucumber (Supplemental Fig. S4). Interestingly, the polymorphism type of *OsUMP1* from R2115 was not in any of those 157 haplotypes. Among the identified SNPs or InDels, 13 were specific for R2115 (Fig. 5A; Supplemental Table S8), suggesting that the *OsUMP1^R2115^* allele was rare in rice populations.

**Figure 5.**
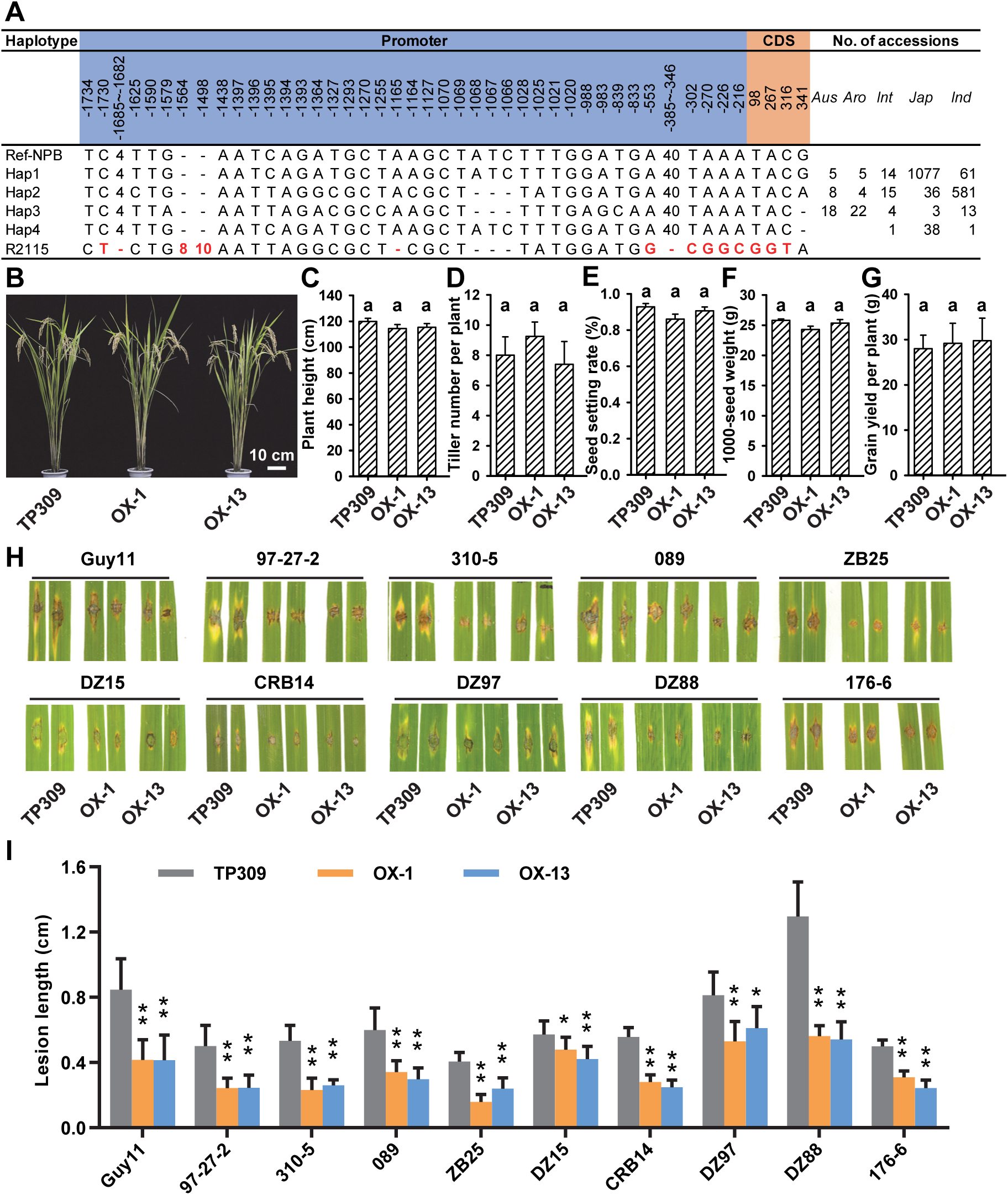
*OsUMP1^R2115^* is a rare superior allele in rice populations. **A,** Polymorphism analysis of *OsUMP1* genomic sequences in rice populations deposited in MBKbase database (http://mbkbase.org/rice). Red letters and numbers indicate R2115-specific SNPs and InDels. **B,** Gross morphology of TP309 and *OsUMP1^R2115^* transgenic lines. **C-G,** Measurement of agronomic traits, including plant height (C), tiller number per plant (D), seed setting rate (E), 1000-seed weight (F), and grain yield per plant (G). Data are means ± SD of five biological replicates. Significance was determined by one-way ANOVA with post hoc Tukey HSD analysis. **H,** Disease phenotype of indicated rice at 5 days post inoculation (dpi) of a diverse panel of *M. oryzae* strains. **I,** Quantification of lesion length in (H). Data are means ± SD of six biological replicates. Significant difference between TP309 and OX lines was determined by Student’s *t*-test (** *P* < 0.01).

To evaluate the potential application value of *OsUMP1^R2115^* in rice breeding, we assessed whether introduction of *OsUMP1^R2115^* caused trade-off between resistance and growth. We measured the main agronomic traits of *OsUMP1^R2115^*-expressing transgenic lines (OX-1 and OX-13) and found no obvious morphology difference of mature rice plants compared to the TP309 control (Fig. 5B). Also, major yield traits were not significantly affected, leading to comparable grain yield to TP309 (Fig. 5C-G).

We further evaluated the resistance spectrum of *OsUMP1^R2115^* lines to a diverse panel of *M. oryzae* isolates possessing different avirulence genes, e.g. CRB14 harboring *Avr-Pita2*, *Avr-Pi9*, and *Avr-Pi11* (Fang et al., 2018), DZ88 harboring *Avr-Pii*, *Avr-Pik*, and *Avr-Piz5* (data not shown). Markedly, *OsUMP1^R2115^* lines developed much smaller lesions than TP309 to all the tested *M. oryzae* isolates (Fig. 5H, I), indicating *OsUMP1^R2115^* confers race-nonspecific resistance to *M. oryzae*. We next tested the resistance of *OsUMP1^R2115^* plants to two other major foliage diseases, including rice sheath blight and bacterial blight. Mycelial clumps of *R. solani* strain AG1-IA or bacterial suspension of the *Xoo* strain P6 were inoculated onto fully expanded leaves of TP309 and *OsUMP1^R2115^* at tillering stage. At 2 dpi, *R. solani* formed significantly smaller lesions in *OsUMP1^R2115^* than in TP309 (Supplemental Fig. S5A). At 14 dpi, *Xoo* developed shorter lesions in the transgenic lines than in TP309 (Supplemental Fig. S5B). These data suggest a role of *OsUMP1^R2115^* in conferring BSR against multiple leaf-infecting pathogens.

We also evaluated the sensitivity of *OsUMP1^R2115^* plants to an emerging floral pathogen *U. virens*. At rice late booting stage, a field *U. virens* isolate PJ52-2-5 was inoculated into the developing panicles of *OsUMP1^R2115^* plants and TP309. At around four-week post inoculation, greenish-black false smut balls were developed from the infected spikelets (Supplemental Fig. S5C). The average number of false smut balls per diseased panicle in *OsUMP1^R2115^* lines (OX-1 and OX-13) was significantly less than that in TP309 (Supplemental Fig. S5D). Similar trends were observed for the diseased spikelet rate and false smut ball weight per diseased panicle (Supplemental Fig. S5E, F). These results suggest a role of *OsUMP1^R2115^* in promoting false smut resistance in rice, which is further supported by high and *U. virens*-inducible expression of *OsUMP1* in spikelets of *OsUMP1^R2115^* lines (Supplemental Fig. S5G). Accordingly, the defense-related genes (*OsKS4*, *OsPR10b*, and *OsNAC4*) were expressed at higher levels in *OsUMP1^R2115^* lines than in TP309 before/after *U. virens* infection (Supplemental Fig. S5G). Elevation of *OsUMP1^R2115^* expression also led to increases of 26S proteasome accumulation and activity, and decreases of POD and CAT activities (Supplemental Fig. S5H, I). Consequently, the abundance of H_2_O_2_ accumulated significantly higher in the spikelets of *OsUMP1^R2115^* lines than in TP309 control, likely accounting for enhanced resistance to *U. virens* in *OsUMP1^R2115^*.

## DISCUSSION

The ubiquitin-proteasome system (UPS) acts as a hub in plant immunity (Ustun et al., 2016; Copeland and Li, 2019). The UPS is composed of ubiquitin activating enzyme (E1)-ubiquitin conjugating enzyme (E2)-ubiquitin protein ligase (E3) cascade and 26S proteasome. The 26S proteasome consists of 20S core protease (CP) and 19S regulatory particle (RP) that are protein complexes with diverse subunits (Schmidt and Finley, 2014; Xu and Xue, 2019). The roles of E3 ligases in plant immunity have been well-documented (see review Copeland and Li, 2019), but few proteasome subunits have been demonstrated to act in plant immune responses to pathogens (Xu and Xue, 2019). In this work, we provide evidence that a proteasome maturation factor *OsUMP1* regulates rice immunity against diverse pathogens, including fungal and bacterial microbes. Since *OsUMP1* contributes to induction of PTI marker genes and accumulation of H_2_O_2_ and confers race-nonspecific resistance to *M. oryzae*, we hypothesize that *OsUMP1* is a new player in plant basal defense.

UMP1 homologs in *Saccharomyces serevisiae* and human are involved in maturation and assembly of 20S CP; upon completion of 20S CP assembly, UMP1 could be rapidly degraded (Ramos et al., 1998; Burri et al., 2000). Plants UMP1 homologs have been rarely identified except in Arabidopsis (Gemperline et al., 2019). Arabidopsis genome harbors three *UMP1* paralogs including *UMP1a*, *UMP1b*, and *UMP1c*, which are involved in proteotoxic, salt, and osmotic stress responses (Gemperline et al., 2019). In rice, we also found three *UMP1* paralogs, which displayed different expression patterns in response to *M. oryzae* infection (Supplemental Fig. S2). Two rice paralogs, *OsUMP1* and Os09g0314900, expressed in a variety-specific manner, while the other one Os02g0800100 was universally expressed in different tested rice varieties, indicating diverse roles of these paralogs. Using transgenic approaches, we further demonstrated that *OsUMP1* contributed to the abundance and activity of 26S proteasome (Fig. 4A, B).

Proteasome is a general machinery for protein degradation. In consistent with this notion, we observed global proteome changes in *OsUMP1*-overexpressing lines with increased 26S proteasome activity (Supplemental Table S6). Nevertheless, through a yet-to-be discovered mechanism, overexpression of *OsUMP1* led to coordinate reduction of a subset of stress-responsive proteins particularly several POD and CAT proteins responsible for H_2_O_2_ degradation, in turn promoting H_2_O_2_ accumulation (Fig. 4). H_2_O_2_ can activate plant immune responses or directly restricts pathogens (Levine et al., 1994; Chamnongpol et al., 1998), suggesting that H_2_O_2_-degrading enzymes POD and CAT are involved in plant immunity. Indeed, the role of POD in disease resistance has been demonstrated by that down-regulation of POD-encoding genes via a natural allele of transcription factor *bsr-d1* leads to enhanced rice resistance against *M. oryzae* (Li et al., 2017). Inhibited expression of CAT-encoding genes can also activate plant defense responses and enhance resistance to mosaic virus in tobacco (Takahashi et al., 1997). Thus, our data indicate a OsUMP1-mediated defense pathway for regulation of POD and CAT protein levels to manipulate H_2_O_2_ homeostasis.

Crops with multi-disease resistance and high yield are highly desirable by breeders and farmers. Although a list of genes associated with multi-disease resistance have been documented, only a limited number of them can balance resistance and yield due to the trade-off between immunity and growth (Li et al., 2020). For instance, wheat *Fhb*7 confers resistance to FHB and crown rot without yield penalty (Wang et al., 2020). Rice *Ideal Plant Architecture 1* (*IPA1*) not only activates yield-related genes to promote grain yield but also enhances rice blast resistance through a phosphorylation regulation mechanism (Wang et al., 2018). Inducible overexpression of *IPA1* enhances resistance to bacterial blight and grain yield (Liu et al., 2019). In our study, the novel *OsUMP1^R2115^* allele confers rice resistance to fungal biotroph *U. virens*, hemibiotroph *M. oryzae*, necrotroph *R. solani* and bacterial hemibiotroph *Xoo* (Fig. 2, 5; Supplemental Fig. S5), representing a new broad-spectrum and multi-disease resistance gene. Furthermore, transgene of *OsUMP1^R2115^* in a susceptible rice accession caused no observable penalties in yield traits (Fig. 5B-G). It is worthy to investigate how *OsUMP1^R2115^* balances multi-disease resistance and yield.

Compared to the coding sequence of *OsUMP1*, there were more variations in its promoter region among rice populations (Fig. 5A), indicating that *OsUMP1* is subjected to dynamic regulations. Then, *OsUMP1* alleles that can balance different agricultural traits would be expected. The *OsUMP1^R2115^* allele balancing BSR and yield traits may be attributed to pathogen-inducible expression of *OsUMP1^R2115^* (Fig. 3B; Supplemental Fig. S5G). Future work is needed to identify the pathogen-responsive cis-elements in the promoter of *OsUMP1^R2115^*. As the *OsUMP1^R2115^* allele is rare in rice populations (Fig. 5A; Supplemental Table S8), it is of great potential to introduce *OsUMP1^R2115^* into other rice varieties. Moreover, identification of *OsUMP1^R2115^* will open a way for dissecting the resistance mechanism of rice against the floral pathogen *U. virens*.

## MATERIALS AND METHODS

### Rice Accessions and Fungi

Rice (*Oryza sativa* L.) accessions used in this study include an elite hybrid rice restorer line Yahui2115 (R2115), a blast-susceptible *japonica* accession Taipei 309 (TP309), a blast-susceptible *japonica* accession Lijiangxin Tuan Heigu (LTH) and three of its derived monogenic resistant lines IRBLz5-CA, IRBLz9-W, and IRBLkm-Ts harboring blast resistance gene *Pi2, Pi9*, and *Pikm*, respectively. Rice plants were maintained under 14 h light/10 h darkness at 27 ± 2°C and 70% relative humidity. *M. oryzae* strains used in this study include Guy11, an eGFP-tagged strain Zhong8-10-14 (GZ8), and a panel of field isolates (176-6, 176-67, 145-8, 147-10, 98-2, 97-27-2, DZ15, DZ88, DZ97, ZB25, 089, 310-5, and CRB14). *M. oryzae* strains were cultured in complete media at 28°C. *U. virens* strain PJ52-2-5 (PJ52), *R. solani* strain AG1-IA, and *Xoo* strain P6 were cultured in potato-sucrose-agar (PSA) media at 28°C.

### Constructs and genetic transformation

To knockout *OsUMP1* in TP309, we applied a CRISPR-Cas9 approach by designing a gene-specific guide RNA with an online software toolkit CRISPR-GE (Xie et al., 2017) and constructing Cas9-*OsUMP1* plasmid with the pRGEB32 binary vector (Addgene, USA) as described previously (Xie et al., 2015). Construct Cas9-*OsUMP1* was genetically transformed into TP309 mediated by *Agrobacterium* strain EHA105. The resultant transgenic plants were tested for genome editing in *OsUMP1* by PCR and DNA sequencing. Primers are listed in Supplemental Table S9.

To generate *OsUMP1*-overexpressing construct, the genomic fragment, including gene region and its 2254-bp upstream promoter sequence, was amplified from R2115 gDNA and cloned into the pCAMBIA1300 vector, leading to a recombinant plasmid p1300-OsUMP1^R2115^. This construct was introduced into TP309 via EHA105-mediated transformation. Hygromycin B was used for screening the positive transgenic lines. The expression levels of *OsUMP1* in transgenic plants were determined by RT-qPCR. Primers are included in Supplemental Table S9.

### Disease Assay

For spray inoculation of *M. oryzae*, four-leaf-stage rice seedlings were challenged with *M. oryzae* spores at a concentration of 5×10^5^ spore ml^-1^ as described previously (Li et al., 2014). For punch inoculation of *M. oryzae*, the second newly expanded leaves from four- to eight-leaf-stage rice plants were wounded and challenged with *M. oryzae* spores at a concentration of 5×10^5^ spore ml^-1^ as described (Li et al., 2017). The disease phenotype was recorded within seven days post inoculation (dpi). Lesion length was measured with ImageJ software (https://imagej.net). Relative *M. oryzae* biomass was quantified as described (Park et al., 2012).

For inoculation of *U. virens*, rice panicles at late booting stages (5-7 days before heading) were injected with mixture of mycelia and conidia (1×10^6^ conidia ml^-1^) as performed previously (Fan et al., 2015). Disease phenotype and the number/weight of false smut balls were recorded at four weeks post inoculation. For inoculation of *R. solani*, fully expanded leaves at tillering stage were inoculated with mycelial clumps following a previous method (Jia et al., 2007). For inoculation of *Xoo*, fully expanded leaves at tillering stage were challenged with bacterial suspension (OD_600_=0.6) by a clipping method (Han et al., 2014).

### RNA-Seq and data analysis

*M. oryzae* spores of mixed strains (176-6, 176-67, 145-8, 147-10, and 98-2) were spray-inoculated onto seedlings of LTH, R2115, IRBLz5-CA, IRBLz9-W, and IRBLkm-Ts. Two biological replicates (each from 3-5 seedlings) of leaves were collected at 0, 12, and 24 hpi, and subjected to RNA extraction, RNA-Seq library construction, and Illumina HiSeq 2500 sequencing at Novogene Technology Co., Ltd. (Beijing, China). Clean reads were mapped to a reference Nipponbare genome (ftp://ftp.ensemblgenomes.org/pub/release-31/plants/fasta/oryza_sativa/dna/) by TopHat2 (Kim et al., 2013). Gene quantity was calculated by HTSeq (Anders et al., 2015) and differentially expressed genes (DEGs) were identified by DESeq2 (Love et al., 2014). All DEGs were clustered by a Hierarchical-clustering method with in-house R scripts.

### RT-qPCR, H_2_O_2_ measurement, and microscopic observation

Total RNA was isolated by TRIzol reagent (Invitrogen) and was reverse transcribed by the PrimeScript^TM^ RT reagent Kit with gDNA Eraser following the manufacturer’s instructions (TaKaRa). RT-qPCR was performed with SYBR Green mix (Bimake). *OsUbi* was served as an internal control for quantification of relative gene expression. Gene-specific primers are presented in Supplemental Table S9.

Rice leaves spray-inoculated with *M. oryzae* were stained with 1 mg ml^-1^ DAB (Sigma) for visualization of H_2_O_2_ accumulation according to a previous report (Li et al., 2014). Photos were taken under a stereomicroscope SZX16 (Olympus) and a Axio Imager A2 (Zeiss). Leaves or spikelets challenged with *M. oryzae* or *U. virens* were sampled for quantification of H_2_O_2_ with the Amplex Red Hydrogen Peroxide/Peroxidase Assay kit (Invitrogen) following the manufacturer’s instructions and a previous report (Li et al., 2019).

To monitor the infection progression of *M. oryzae* in rice, spores (1×10^5^ spore ml^-1^) of an eGFP-tagged strain GZ8 were inoculated into rice sheaths from six-week-old plants. The epidermal layer from the inoculated leaf sheaths was observed under a Laser Scanning Confocal Microscope (Nikon A1) at 18 and 40 hpi. The infection phase of over 150 spores was recorded at each time point for each rice line.

### Proteomic analysis

At four-leaf stage, three biological replicates of leaves (each from 10 individual plants) of TP309 and *OsUMP1*-overexpressing line OX-1 were sampled. Protein extraction, digestion, and iTRAQ labeling were performed as previously described (Fan et al., 2011). TP309 replicated samples were labeled with 113, 114, and 115, OX-1 replicated samples were labeled with 116, 117, and 118. After labeling, the samples were mixed and subjected to fractionation by strong cation exchange (SCX) chromatography using the AKTA Purifier system (GE Healthcare). Fractions were desalted on C18 Cartridges (Empore™ SPE Cartridges C18, bed I.D. 7 mm, volume 3 ml, Sigma), and reconstituted in 0.1% (v/v) trifluoroacetic acid after vacuum centrifugation. Subsequently, nanoLC-MS/MS analysis was performed on a Q Exactive mass spectrometer coupled with Easy nLC (Proxeon Biosystems, now Thermo Fisher Scientific). MS/MS spectra were analyzed with Proteome Discoverer 1.3 (Thermo Electron) coupled with MASCOT engine (Matrix Science) against Uniprot database (https://www.uniprot.org/), with the built-in settings of the software. Above iTRAQ experiments were conducted at the Applied Protein Technology (Shanghai). Note that one biological replicate of TP309 was of low quality and thus was excluded from analysis of differentially accumulated proteins. Gene ontology enrichment analysis of differentially accumulated proteins was performed with KOBAS 2.0 web server (Xie et al., 2011).

### ELISA

The amounts of 26S proteasome were determined with the Plant 26S proteasome (26S PSM) ELISA Kit (JKBIO, Shanghai) following the manufacturer’s instructions. In brief, 50 mg fresh leaves powdered in liquid nitrogen and homogenized with 150 μl of Kit-provided PBS. The extracts were centrifuged at 5000 rpm for 10 min at 4°C, and the supernatants were subjected to measurements with a 96-well Thermo Scientific Microplate Reader at 450 nm. The amounts of POD and CAT were measured with the Plant peroxidase (POD) ELISA Kit (mibio, Shanghai,) and Plant Catalase (CAT) ELISA Kit (mibio, Shanghai), respectively, following the manufacturer’s instructions.

### Measurement of proteasome, POD, and CAT activity

Proteasome activity was determined as described (Üstün and Börnke, 2017). Briefly, 200 mg of rice leaves were collected and ground in 200 μl extraction buffer (50 mM pH 7.2 HEPES-KOH, 250 mM Suc, 2 mM dithiothreitol, and 2 mM ATP). Homogenized samples were centrifuged, and the supernatant was collected and adjusted to 1 mg ml^-1^. Thirty micrograms of isolated protein were added into the wells of a black-walled 96-well plate and subsequently mixed with 220 μl assay buffer (100 mM pH 7.8 HEPES-KOH, 2 mM ATP, 10 mM KCl, and 5 mM MgCl_2_). Proteasome substrate Suc-LLVY-AMC (Sigma) was added into the wells and incubated for 15 min at 30 °C. The release of amino-methyl-coumarin was recorded with a Thermo Scientific Microplate Reader (excitation at 360 nm and emission at 460 nm) every 1.5 min over a period of 120 min.

The enzyme activities of POD and CAT were measured with the POD Activity Assay Kit and CAT Activity Assay Kit (Solarbio, Beijing) following the manufacturer’s directions. Briefly, 50 mg of rice leaves were collected and ground in 100 μl of Kit-provided PBS. After centrifugation, the supernatants were used for enzyme activity measurement with a 96-well Thermo Scientific Microplate Reader at specific wavelengths (470 nm for POD and 240 nm for CAT).

### Sequence analysis

The genomic sequences of *OsUMP1* for each haplotype were retrieved from the database of MBKbase-rice (http://mbkbase.org/rice). The genomic sequence of *OsUMP1^R2115^* was amplified from R2115 and confirmed by sequencing with multiple clones. Sequence alignment was conducted with the Muscle method and phylogenetic tree was constructed with the Neighbor-Joining method in MEGA5.1 (Tamura et al., 2011). Corresponding protein sequences were deduced by online software (https://web.expasy.org/translate).

## Supplemental Data

The following materials are available in the online version of this article.

**Supplemental Figure S1.** Expression patterns for subcluster 5-10 of differentially expressed genes.

**Supplemental Figure S2.** Expression patterns of *OsUMP1* homologs in rice responsive to *Magnaporthe oryzae* infection.

**Supplemental Figure S3.** *OsUMP1* contributes to rice resistance to *Magnaporthe oryzae*.

**Supplemental Figure S4.** Sequence alignment and phylogenetic analysis of OsUMP1.

**Supplemental Figure S5.** The *OsUMP1^R2115^* allele confers resistance to foliar and floral pathogens.

**Supplemental Table S1.** Mapping results of RNA-Seq reads.

**Supplemental Table S2.** The complete list of differentially expressed genes from rice in response to *Magnaporthe oryzae* infection (the data represent normalized FPKM).

**Supplemental Table S3.** GO enrichment analysis for genes in Subcluster 3.

**Supplemental Table S4.** KEGG enrichment analysis for genes in Subcluster 3.

**Supplemental Table S5.** Expression data of subcluster 3 genes involved in programmed cell death and proteasome.

**Supplemental Table S6.** Proteomic analysis of OsUMP1R2115 transgenic plants compared with wild-type TP309.

**Supplemental Table S7.** GO enrichment analysis for differentially accumulated proteins in *OsUMP1*-overexpressing line OX-1 compared with TP309.

**Supplemental Table S8.** Natural variations of *OsUMP1* in rice populations.

**Supplemental Table S9.** Primers used in this study.

## ACKNOWLEDGEMENTS

We thank Dr. Cailin Lei (Chinese Academy of Agricultural Sciences) for providing rice lines IRBLZ5-CA, IRBLZ9-W, and IRBLKM-TS; Lihuang Zhu (Chinese Academy of Sciences) for providing the eGFP-tagged *M. oryzae* strain GZ8; Wensheng Zhao (China Agricultural University) for providing *M. oryzae* strain CRB14; Xuewei Chen and Ai-Ping Zheng (Sichuan Agricultural University) for providing *Xoo* strain P6 and *R. solani* strain AG1-IA, respectively; Yangwen Qian (Hangzhou Biogle Co., Ltd.) for assistance with rice genetic transformation.

## Author Contributions

J.F. and W.-M.W. designed the study and wrote the manuscript. X.-H.H., J.F., J.-L.W., S.S., J.-X.H., J.L., H.W., Y.Z., G.-B.L., J.-H.Z., D.-Q.L. M.P., Z.-X.Z., S.-X.Z., J.-W.Z., Y.-Y.H., Y.L., and F.H. performed the experiments. J.F., W.-M.W., X.-H.H., J.X., Y.-L.L., and J.-L.W. analyzed the data. The authors declare no conflict of interest.

## Funding Information

This work was supported by the grants from the National Natural Science Foundation of China (U19A2033, 31672090 and 31430072 to W.-M.W., 31772241 and 32072503 to J.F.), and the Sichuan Applied Fundamental Research Foundation (2020YJ0332 to W.-M.W., and 2020YJ0145 to J.F.).

## Supplemental figures

**Supplemental Figure S1.**
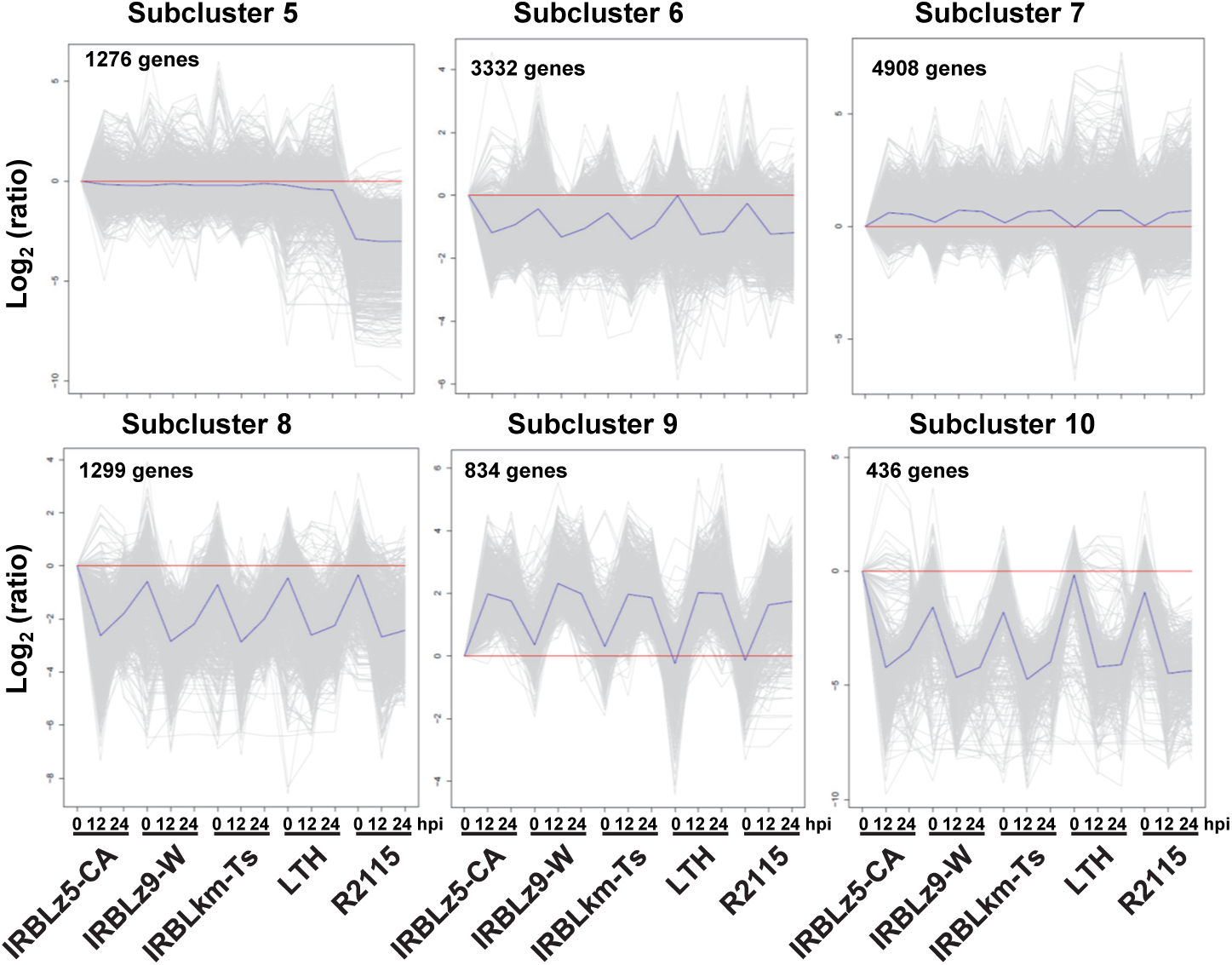
Expression patterns for Subcluster 5-10 of differentially expressed genes. The gray lines in each subgraph represent the relative expression level of genes. The blue lines represent the average relative expression level of all genes in this cluster.

**Supplemental Figure S2.**
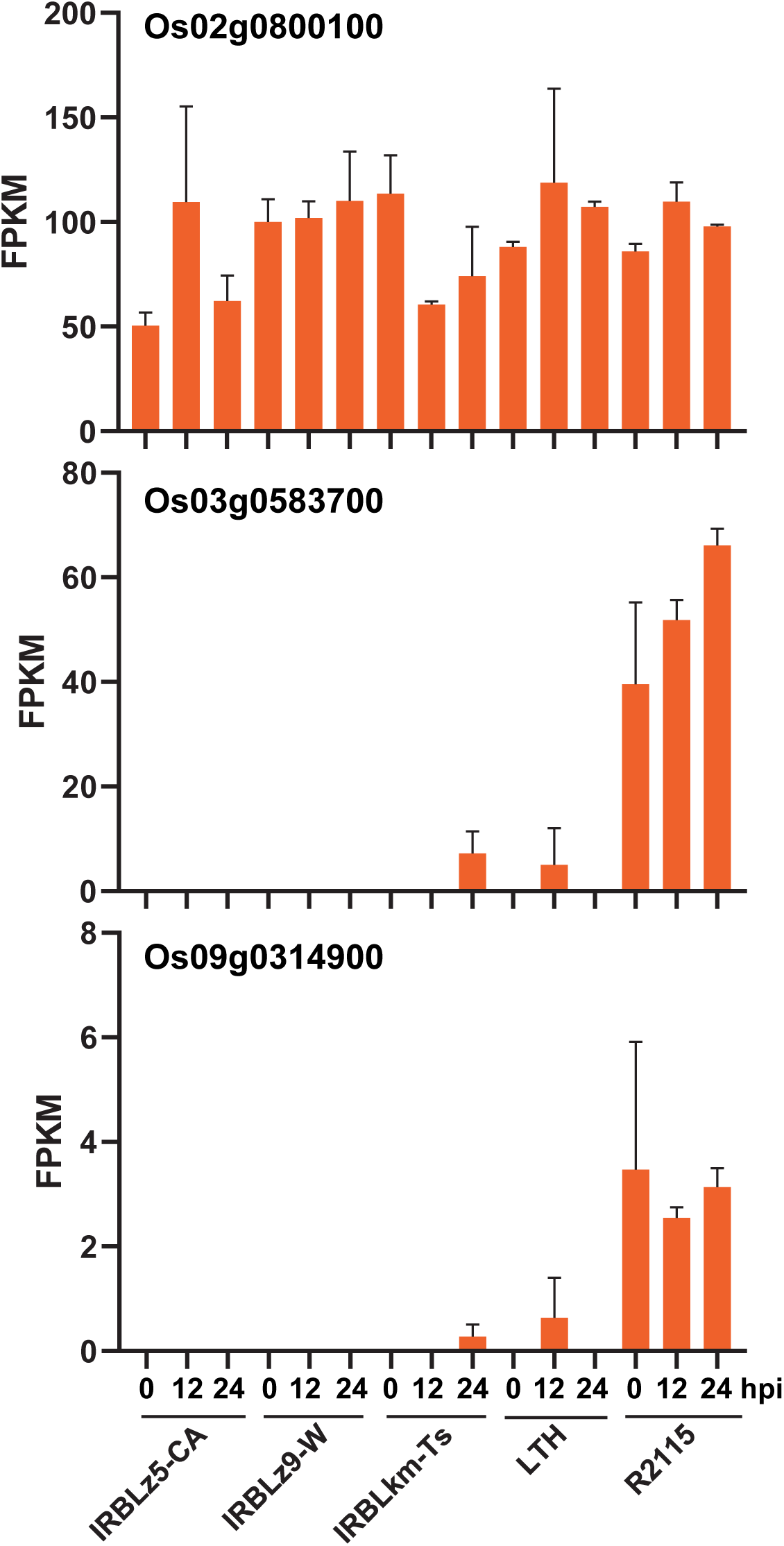
Expression patterns of *OsUMP1* homologs in rice responsive to *Magnaporthe oryzae* infection. The FPKM data were retrieved from the transcriptome data sets in this study.

**Supplemental Figure S3.**
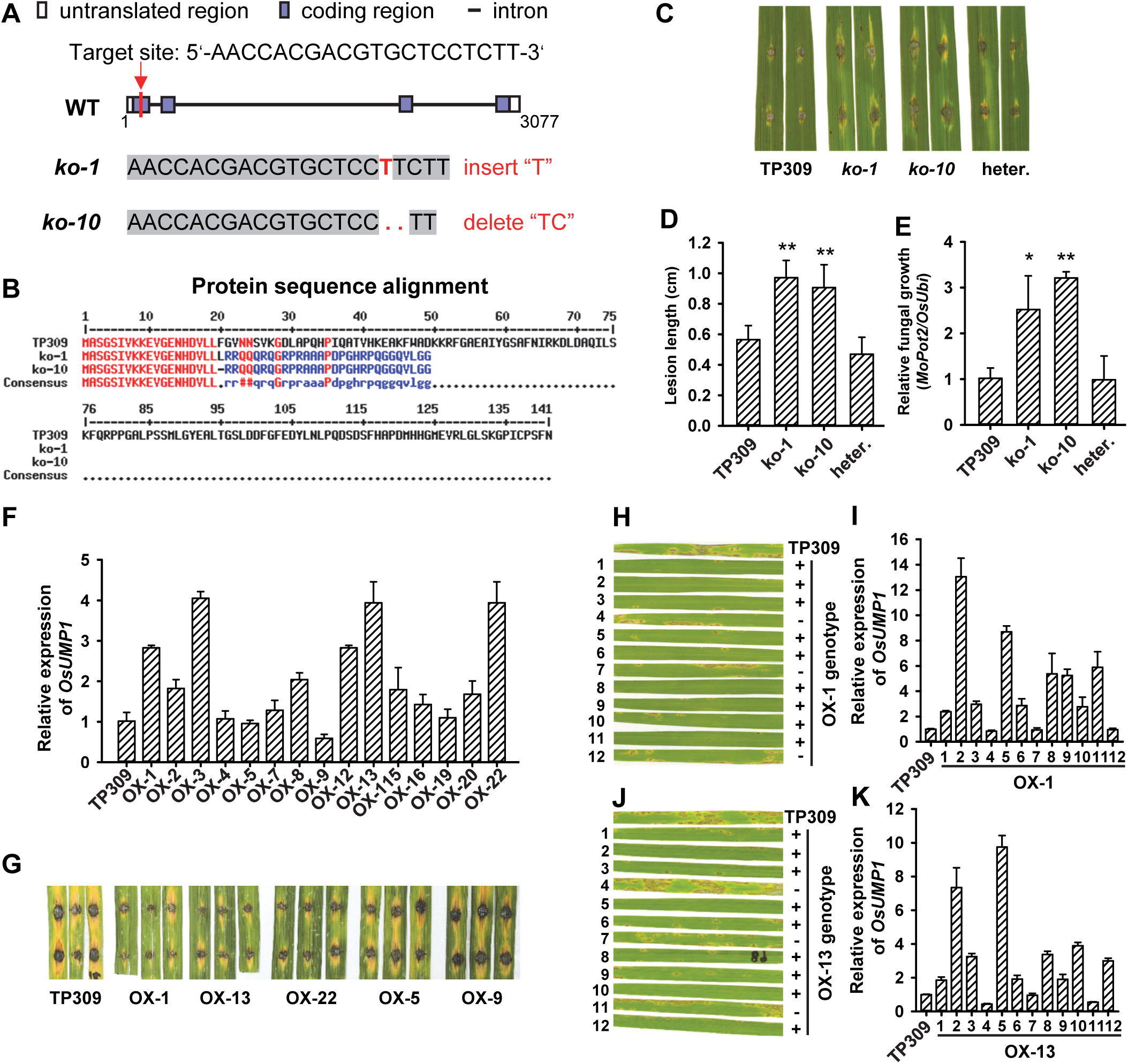
O*s*UMP1 contributes to rice resistance to *Magnaporthe oryzae*. **A,** Schematic diagram of *OsUMP1* and mutation types of CRISPR-Cas9-edited plants. **B,** Protein sequence alignment of OsUMP1 in TP309 and two mutants *ko-1* and *ko-10*. **C,** Disease phenotype of *OsUMP1* knockout lines (T0 generation) at 5 days post inoculation (dpi) of *M. oryzae* GZ8. heter., heterozygous mutant line. **D** and **E,** Quantification of lesion length (C) and relative fungal growth (D). Values are means ± SD of six replicates. Asterisks indicate significant differences between TP309 and knockout lines as determined by Student’s *t*-test (** *P* < 0.01). **F,** Expression of *OsUMP1* in multiple transgenic lines (T0 generation). Os*Ubi* was used as the reference gene and the expression of *OsUMP1* in TP309 was set as the control. **G,** Disease phenotype of multiple *OsUMP1* transgenic lines (T0 generation) at 6 dpi of GZ8. **H**-**K,** Disease phenotype of OX-1 and OX-13 segregants inoculated with *M. oryzae* GZ8. The expression levels of *OsUMP1* in these progenies co-segregated with their resistance to *M. oryzae*. RT-qPCR analysis was performed with Os*Ubi* as the reference gene and the expression of *OsUMP1* in TP309 as the control. Values are means ± SD of three replicates.

**Supplemental Figure S4.**
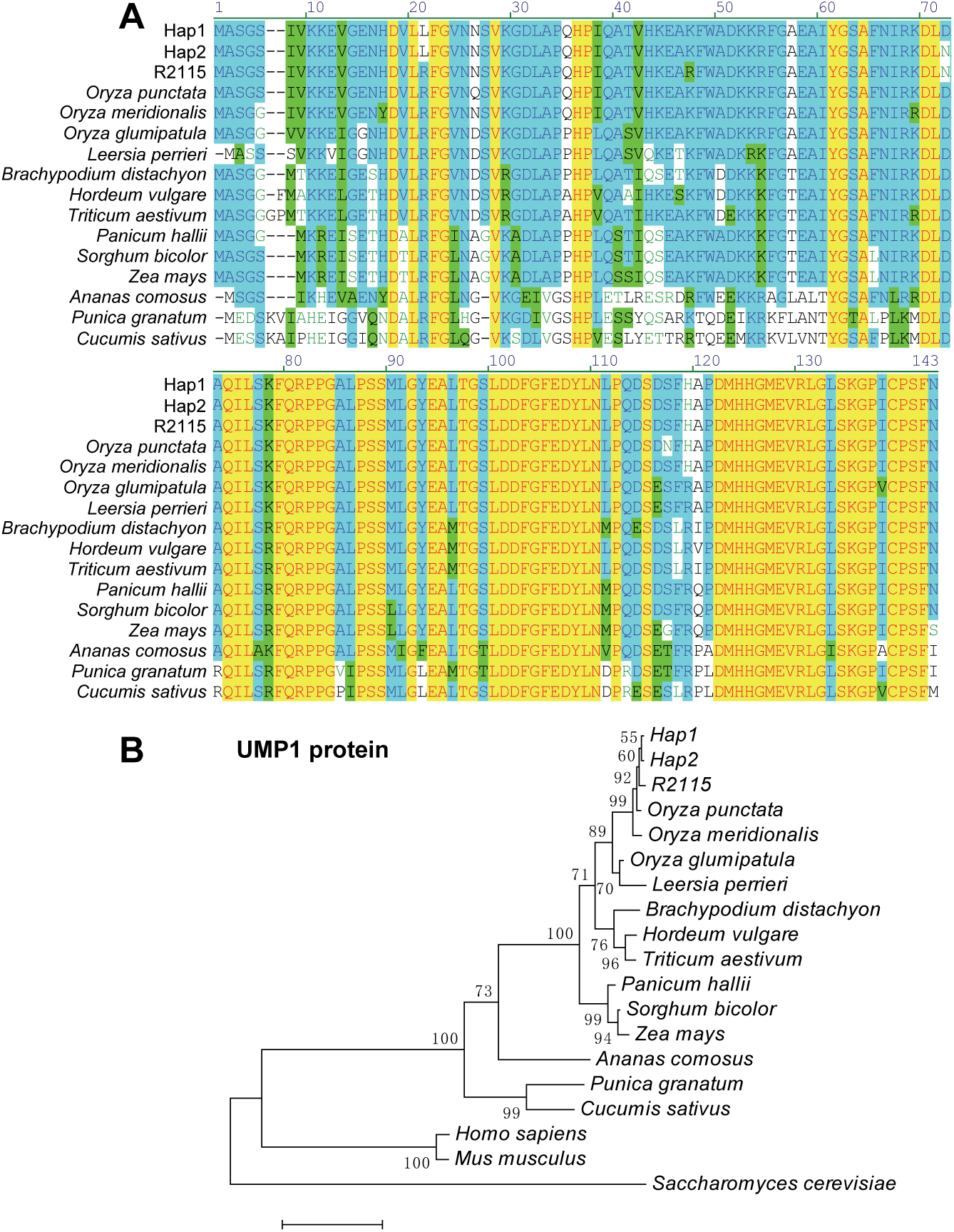
Sequence alignment and phylogenetic analysis of OsUMP1. **A,** Protein sequences of OsUMP1 homologs in different plant species were downloaded from the Uniprot database (https://www.uniprot.org/), and aligned using vector NTI software (version 11.5.1). **B,** Phylogenetic tree was generated with MEGA5.1 using the Muscle method for sequence alignment and the Neighbor-Joining method for tree construction. The accession numbers for OsUMP1 homologs are as below: *Oryza punctata* (A0A0E0KFV5), *Oryza meridionalis* (A0A0E0D398), *Oryza glumipatula* (A0A0D9YZR5), *Leersia perrieri* (A0A0D9VM90), *Hordeum vulgare* (F2CQM0), *Triticum aestivum* (A0A3B6QNA6), *Brachypodium distachyon* (I1IDJ7), *Panicum hallii* (A0A2S3GU64), *Sorghum bicolor* (A0A194YTW9), *Zea mays* (B4FLC7), *Ananas comosus* (A0A199VSL7), *Punica granatum* (A0A218XIA6), and *Cucumis sativus* (A0A0A0KUK6). OsUMP1 homologs in *Homo sapiens* (Q9Y244), *Mus musculus* (Q9CQT5), and *Saccharomyces cerevisiae* (P38293) were included as outgroups for phylogenetic analysis.

**Supplemental Figure S5.**
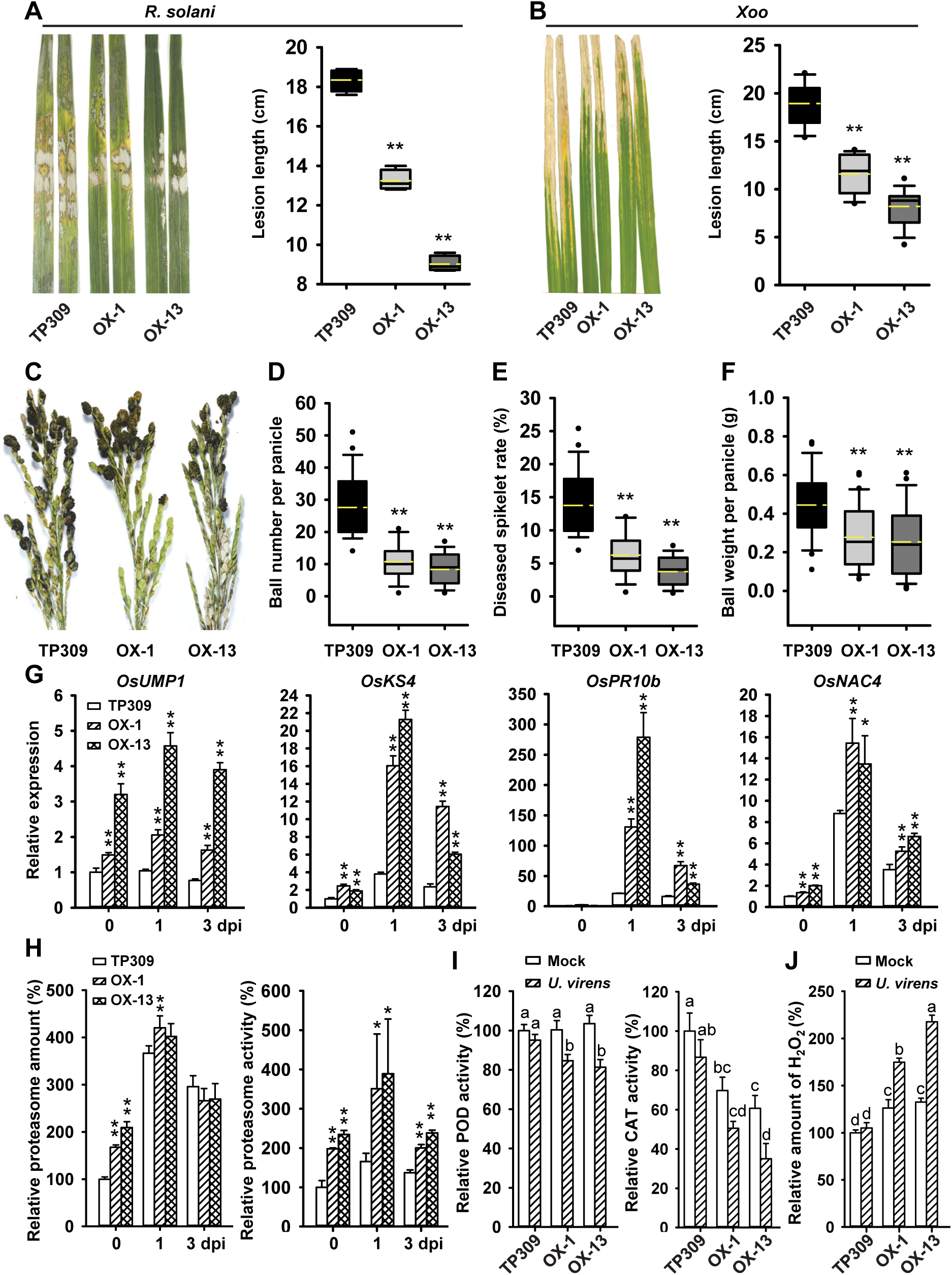
The *OsUMP1^R2115^* allele confers resistance to foliar and floral pathogens. **A,** Disease phenotype and lesion length of indicated rice lines at two days post inoculation (dpi) of *Rhizoctonia solani* AG1-IA. Data are box-plotted (n > 10). Significant difference between TP309 and OX lines was determined by Student’s *t*-test (** *P* < 0.01). **B,** Disease phenotype and lesion length of indicated rice lines at 14 dpi of *Xanthomonas oryzae* pv. *oryzae* P6. Data are box-plotted (n > 20). Significant difference between TP309 and OX lines was determined by Student’s *t*-test (** *P* < 0.01). **C,** Disease phenotype of indicated rice lines at 28 dpi of *Ustilaginoidea virens* PJ52. **D-F,** Statistical analysis of false smut ball number per diseased panicle (D), diseased spikelet rate per panicle (E), and false smut ball weight per diseased panicle (F). Data are box-plotted (n > 30). Significant difference between TP309 and OX lines was determined by Student’s *t*-test (* *P* < 0.05, ** *P* < 0.01). **G,** Expression analysis of *OsUMP1* and defense-related genes in spikelets of indicated rice lines after inoculation of PJ52. *OsUbi* was used as the reference gene. The expression of indicated genes in TP309-0 dpi was set as the control. Data are means ± SD of three biological replicates. Significant difference between TP309 and OX lines was determined by Student’s *t*-test (** *P* < 0.01). **H,** The 26S proteasome amount and activity in spikelets of indicated rice lines after inoculation of PJ52. Data are means ± SD of three biological replicates. Significant difference between TP309 and OX lines was determined by Student’s *t*-test (* *P* < 0.05, ** *P* < 0.01). **I and J,** Enzyme activities of POD and CAT (I) and H_2_O_2_ quantification in spikelets of indicated rice lines at 1 dpi of PJ52. Values represent means ± SD of three biological replicates. Different letters indicate significant differences as determined by one-way ANOVA with post hoc Tukey HSD analysis.

## Notes

### Competing Interest Statement

The authors have declared no competing interest.

